# Albendazole specifically disrupts microtubules and protein turnover in the tegument of the cestode *Mesocestoides corti*

**DOI:** 10.1101/2025.03.04.641463

**Authors:** Inés Guarnaschelli, Uriel Koziol

## Abstract

Parasitic flatworms, such as cestodes and trematodes, are covered by a syncytial tissue known as the tegument. It consists of a superficial band of cytoplasm (the distal tegument) that is connected by cytoplasmic bridges to multiple cell bodies (cytons) that lay beneath the basal lamina and contain the nuclei. We characterized the cytoskeleton of the tegument of the model cestode *Mesocestoides corti* and determined the effects of albendazole and albendazole sulfoxide on its organization. These anthelmintics are known to target beta-tubulin in helminths, and their effects have been extensively studied in nematodes, but the specific cells and tissues that are affected are not well understood in parasitic flatworms. Using antibodies that detect different tubulin subunits and post-translational modifications, we show that microtubules in the distal tegument have a unique organization, with bouquets of microtubules radiating from the cytoplasmic bridges, suggesting a role in intracellular traffic. In contrast, actin filaments were largely absent from the distal tegument. The microtubules of the tegument were specifically sensitive to low, chemotherapeutically relevant concentrations of albendazole and albendazole sulfoxide. This was correlated with the accumulation of secretory material in the cytons, and low concentrations of albendazole strongly reduced the incorporation of newly synthesized proteins in the distal tegument, as determined by metabolic labeling. Unexpectedly, albendazole also induced a global decrease in protein synthesis, which was independent of the activation of the unfolded protein response. Our work identifies the tegument as a sensitive target of benzimidazoles in cestodes and indicates that translational inhibition may contribute to the anthelmintic effect of benzimidazoles.

**Author Summary:** Infections caused by cestodes, such as *Echinococcus* spp. and *Taenia* spp., pose a large burden on global human health. These parasites are covered by a very particular epidermis, a syncytial tegument, which is the only contact surface between the parasite and the host. The only chemotherapy available for these infections are benzimidazoles, such as albendazole. From work in other helminths, it is well known that beta-tubulin is the molecular target of benzimidazoles, however, the affected cells and tissues are not known in flatworms. Here, we use the model cestode *Mesocestoides corti* to address some basic features of the tegument such as the organization of the cytoskeleton, and the effects of therapeutically relevant concentrations of albendazole in this tissue. We found that microtubules are abundant and highly organized in the tegument, and are specifically sensitive to albendazole, in comparison with other cells and tissues. Albendazole reduced the incorporation of new proteins in the tegument and produced an unexpected overall decrease in protein synthesis, which may also be of relevance for chemotherapy. Our results indicate that the tegument is a sensitive cellular target of benzimidazoles.

## Introduction

Parasitic flatworms, such as cestodes (tapeworms) and trematodes (flukes) pose a large burden on global human health, and have a strong impact on livestock productivity [1,2]. One common characteristic of all these parasites is that they lack a typical epidermis, and their body wall is covered instead by a highly specialized syncytial tegument, also known as neodermis [3]. The tegument comprises a superficial band of continuous cytoplasm (known as the distal tegument) that is connected by thin cytoplasmic bridges to individual nucleated cell bodies (cytons) lying beneath the basal lamina (Fig 1A). The surface of the distal tegument is the main site of contact between parasite and host, and shows diverse specializations in different groups, including microvilli-like projections known as microtriches in cestodes. The tegument is a very active tissue, with continuous protein turnover and secretion from the tegumental surface. Classical experiments have shown that protein synthesis is largely absent in the distal tegument. New proteins are continuously synthesized within the cytons (where the endoplasmic reticulum and Golgi complex reside), and transported to the tegumental surface [4,5,6]. Despite its relevance in the infection process, the unique cellular mechanisms required for tegumental turnover and secretion are essentially unknown.

**Fig 1.**
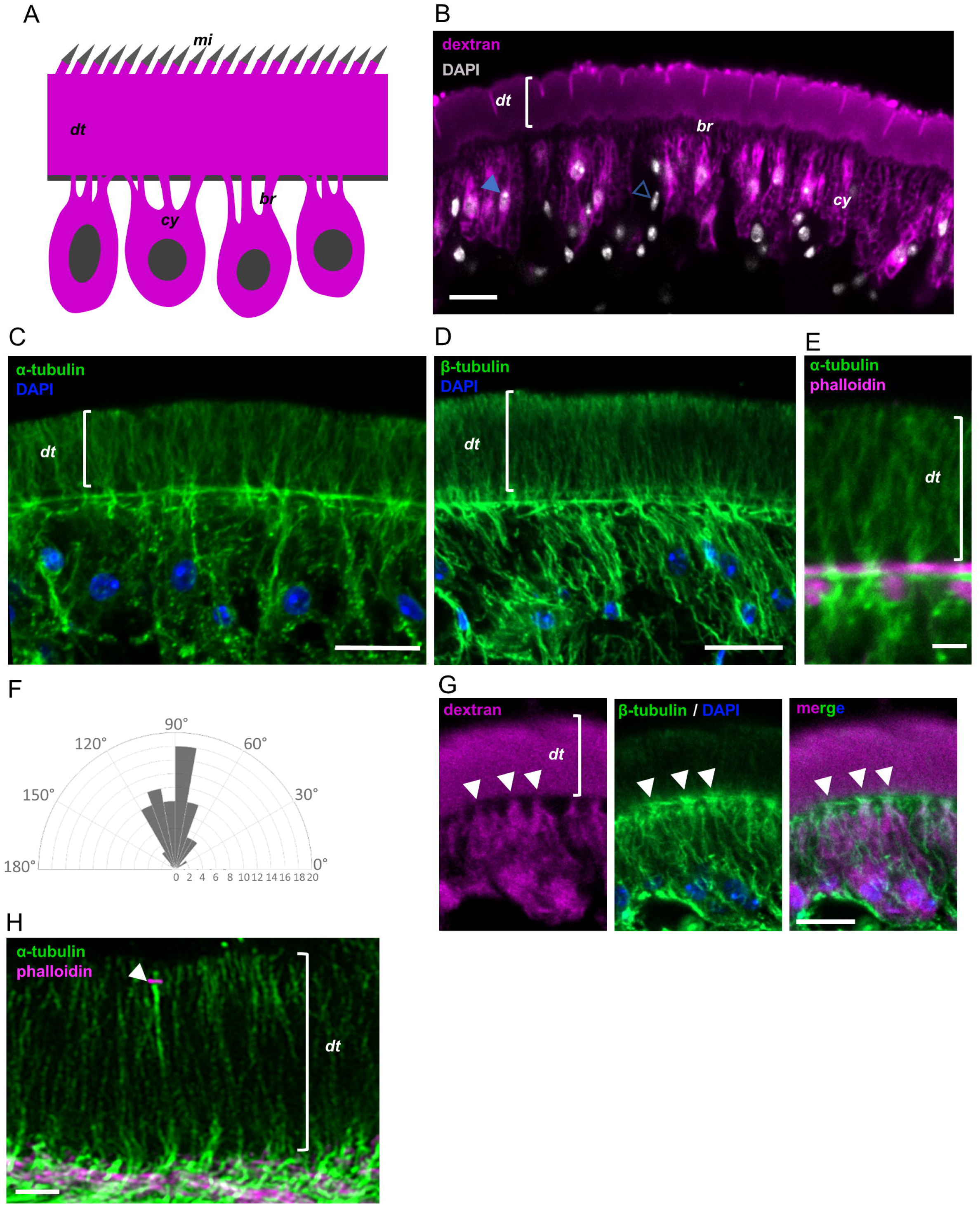
Microtubule organization in the tegument of *M. corti*. (A) Schematic drawing depicting the tegument in cestodes. (B) Tegument labeling with fluorescent dextran. Filled arrowhead points to a dextran-labeled cyton, open arrowhead points to a non-tegumental nucleus in the subtegumental region. The labeling experiment was repeated three times, using different batches of larvae. (C) Detection of alpha-tubulin by immunofluorescence in the tegument. This experiment was repeated three times, using different batches of larvae. (D) Detection of beta-tubulin by immunofluorescence in the tegument. This experiment was repeated three times, using different batches of larvae. (E) Detail of the bouquet-like distribution of microtubules. Phalloidin staining shows muscle fibers adjacent to the basal region of the distal tegument. (F) Wind rose histogram of the angles of orientation of microtubules relative to the basal region of the tegument. The x axis indicates the number of measured angles for each orientation. (G) Co-labeling of the tegument with fluorescent dextran and of microtubules by immunofluorescence, showing that the latter are especially abundant in the cytoplasmic bridges (arrowheads). This experiment was repeated twice, using different batches of larvae. (H) Beta-tubulin distribution imaged with an Airyscan detector. Co-labeling with fluorescent phalloidin shows F-actin associated to the junction between a sensory nerve ending and the distal tegument (arrowhead). This experiment was performed once. mi, microtriches; dt, distal tegument; cy, cytons; br, cytoplasmic bridges. Scale bars: B, C, D and G: 10 μm; E and H: 2 μm.

In classical epithelia from other animals, cells are polarized, particularly regarding the organization of the cytoskeleton. Parallel microtubule bundles with apico-basal orientation are nucleated from their minus ends at the apical cortex, and these are important for polarized vesicular transport and exocytosis at the apical and basolateral domains of the plasma membrane [7,8,9]. In parasitic flatworms, microtubules have been shown to be abundant in tegumental cytons and cytoplasmic bridges by means of electron microscopy, but their overall arrangement and orientation is not well understood [10]. Their arrangement is particularly unclear in the highly crowded distal tegument, which is packed with secretory vesicles. In the cestode *Hymenolepis diminuta*, it has been shown that high concentrations of colchicine, a drug that inhibits microtubule polymerization, can impair incorporation of new proteins into the distal tegument [11,12]. These classical experiments suggest an important role of microtubules in the transport of new proteins from the cytons to the distal tegument.

Benzimidazoles are a large class of anthelmintic drugs, which are active towards many different species of parasitic flatworms and nematodes [13]. In nematodes, there is extensive evidence supporting the identification of beta-tubulin as the molecular target of benzimidazoles, and many parasitic nematode species, particularly gastrointestinal parasites, are highly susceptible to these drugs [14]. Studies in the model nematode species *Caenorhabditis elegans* have identified one beta-tubulin gene, *ben-1*, that is solely responsible for causing benzimidazole susceptibility [15, 16]. Furthermore, genetic resistance to benzimidazoles has been mapped to mutant beta-tubulin alleles in veterinary gastrointestinal nematodes such as *Haemonchus contortus* and *Ancylostoma caninum* [17,18]. At the cellular level, there is evidence of extensive damage to the intestine of gastrointestinal nematodes treated with benzimidazoles [19, 20], whereas in *C. elegans* the target tissue has been shown to be the nervous system [21].

In the case of parasitic flatworms, benzimidazoles (in particular albendazole; ABZ) are the drug of choice for chemotherapeutic treatment against larvae of the cestodes *Echinococcus granulosus* and *Echinococcus multilocularis* (causative agents of cystic and alveolar echinococcosis, respectively), and are also used for the treatment of infections with *Taenia solium* larvae (cysticercosis) [22,23,24,25]. However, the potency of benzimidazoles and their affinity for tubulin are paradoxically low in these parasites [13]. Both ABZ and albendazole sulfoxide (ABZSO), the main active metabolite resulting from the conversion of ABZ in the liver [26], have similar but low potencies *in vitro* against *Echinococcus* and *Taenia* larvae [27,28,29,30,31,32]. Under *in vitro* conditions, parasite damage and death typically require long incubations with ABZ or ABZSO at concentrations of 10 to 40 µM [28,33], one to two orders of magnitude higher than the maximal concentrations found *in vivo* in the plasma of patients (typically around 1 µM) [34,35,36,37]. Furthermore, it is not clear which tissues are affected by benzimidazoles in parasitic flatworms. Unlike nematodes, cestodes lack a digestive system, and the nervous system is reduced to a minimal nerve net in *E. multilocularis* larvae [38]. On the other hand, the tegument has been shown to be affected by high concentrations of benzimidazoles in different parasitic flatworms [39,40,41,33], including pioneer reports that showed depletion of microtubules after mebendazole treatment in cestodes [42,43]. Furthermore, treatment with ABZ also resulted in the accumulation of secretory vesicles in the tegumental cytons of different parasitic flatworms [41]. Thus, the available evidence suggests that tegumental microtubules and intracellular transport may be affected by these drugs.

In this work, we have used the model cestode *Mesocestoides corti* [44] to describe in detail the organization of the cytoskeleton of the tegument, and the effects on this tissue of chemotherapeutically relevant concentrations of ABZ. Our work identifies the tegument as a sensitive target of ABZ in cestodes, and indicates that translational inhibition may contribute to the anthelmintic effect of benzimidazoles.

## Results

### General organization of the tegument of *M. corti*

The tegument of the larva of *M. corti* has been previously described by electron microscopy [45]. The distal tegument has different types of microtriches [46] and varies in thickness along its length, from approximately 5 µm in the anterior end (the scolex) to 11 µm in the posterior end.

The tegumental cytons are known to be interspersed with other cell types in the region beneath the basal lamina (the sub-tegumental region), including muscle cells of the sub-tegumental muscle layer [45]. In order to identify the tegumental cytons in the sub-tegumental region, we used a method for labeling all of the tegument, including the cytons, by introducing fluorescently labeled dextran [47].

Labeling of the tegument in *M. corti* allowed us to clearly identify the tegumental cytons, together with numerous cytoplasmic bridges that connect each of them to the distal tegument, traversing the subtegumental muscle layer in between the muscle fibers (Fig 1B, Fig S1A). Tegumental cytons represent 62% of all nuclei in the subtegumental region (n=204 nuclei counted from 4 labeled worms). The cytons are more abundant in the anterior regions (2.4 x 10^4^ cytons / mm^2^ tegument surface), whereas in the posterior region, in which cell density is lower, tegumental cytons are more sparsely distributed (1.4 x 10^4^ cytons / mm^2^).

### Distribution of microtubules and tubulin post-translational modifications in the tegument

We analyzed the distribution of microtubules by immunofluorescence with antibodies for alpha and beta-tubulin, with essentially identical results (Fig 1C-E). In the distal tegument, microtubules are highly organized as bouquets that radiate from the basal region. On average, these microtubules are perpendicular to the apical surface of the tegument (89.6 ° +/- 21.2 °, standard deviation) (Fig 1F). Co-detection of tubulin by immunofluorescence together with labeling of the tegument using fluorescent dextran demonstrated that microtubules are highly abundant in the cytoplasmic bridges (Fig 1G), and the microtubule bouquets in the distal tegument originate at the connections with these bridges. In the tegumental cytons, microtubules are relatively abundant, but lack a clear overall organization. The distribution of microtubules was further confirmed by imaging experiments at higher resolution, using an Airyscan detector system (Fig 1H).

We also analyzed the distribution of some post-translational modifications of tubulin that can be associated with the stability of microtubules [48]. Tyrosinated tubulin, a form of tubulin that is more common in recently polymerized microtubules, was abundant in the microtubules of the tegument, suggesting that these are highly dynamic (Fig 2A). In contrast, acetylated tubulin, which is associated with stabilized microtubules, was undetectable in the distal tegument (Fig 2B). Interestingly, acetylated tubulin was abundant in the processes of different types of sensory receptors of nerve cells that traverse the distal tegument [46] (these are the only direct cell-to-cell contacts of the distal tegument, and represent the only lateral regions of its plasma membrane) (Fig 2B, S1B,C).

**Fig 2.**
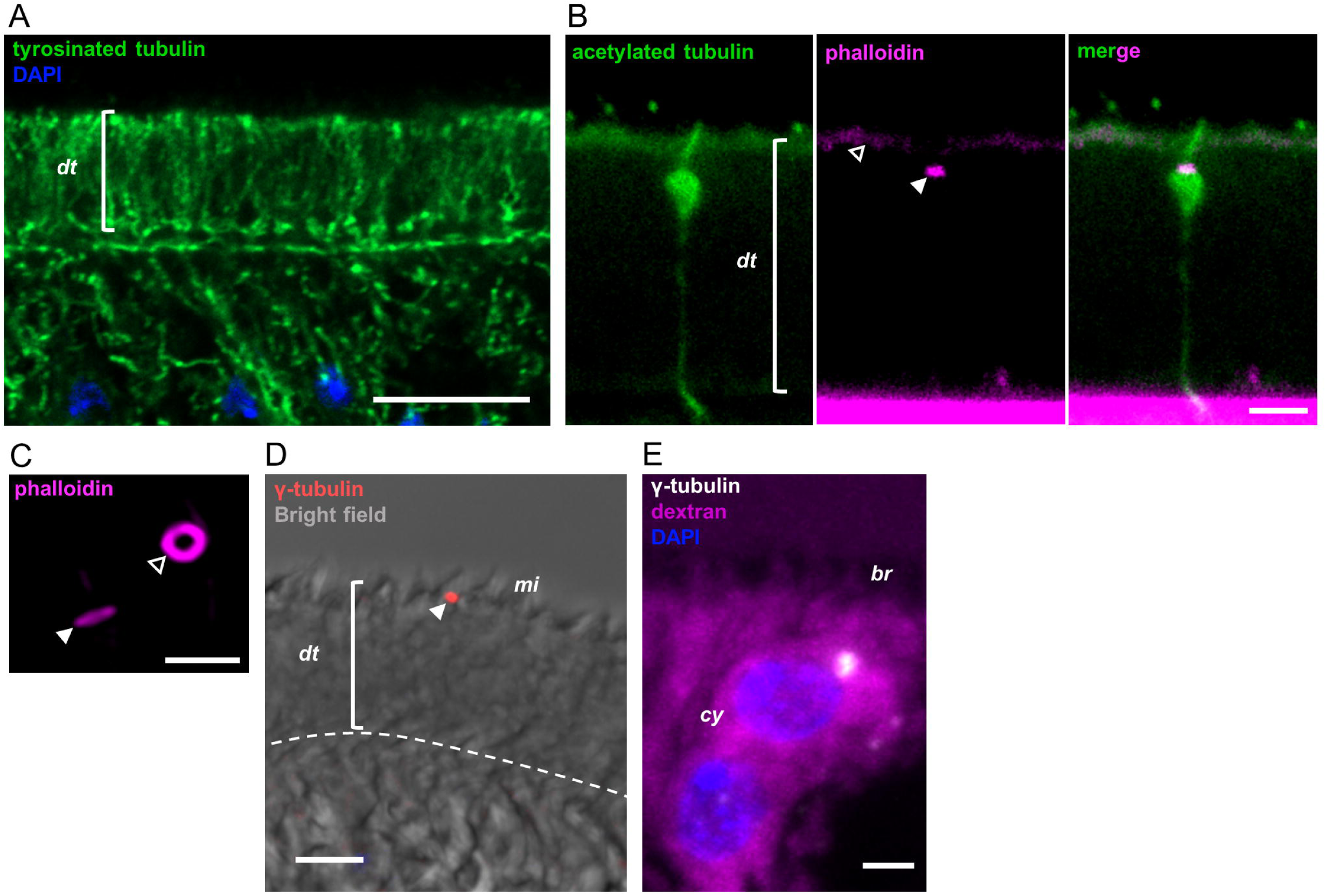
Tubulin modifications and gamma-tubulin distribution in the tegument. (A) Detection of tyrosinated tubulin by immunofluorescence. This experiment was repeated twice using different batches of larvae. (B) Detection of acetylated tubulin by immunofluorescence. Co-labeling with phalloidin shows the actin ring associated to the junction between the sensory nerve ending and the distal tegument (filled arrowhead), as well as actin associated with microtriches (open arrowhead). This experiment was repeated twice using different batches of larvae. (C) Actin rings associated to sensory nerve endings imaged with an Airyscan detector, in lateral (filled arrowhead) and *en face* (open arrowhead) orientations. This experiment was performed once. (D) Detection of gamma-tubulin in the distal tegument, the arrowhead points to a gamma-tubulin speck near the microtriches. This experiment was performed twice. (E) Detection of gamma-tubulin in tegumental cytons (labeled with fluorescent dextran). This experiment was performed once. dt, distal tegument; mi, microtriches; br, cytoplasmic bridges; cy, cytons. Scale bars: A: 10 μm; B, C and E: 2 μm; D: 5 μm.

Actin filaments, labeled with fluorescent phalloidin, were undetectable in the distal tegument with the exception of weak staining of the surface microtriches, and strong staining of actin rings in the cell contacts with the sensory receptors (Fig 2B,C, S1B,C). These probably correspond to the position of septate junctions [49], which have been described between the distal tegument and the sensory receptors [46], and which have been shown to be rich in actin filaments in other species [50].

In order to locate putative microtubule organization centers (MTOCs) within the tegument, we analyzed the localization of gamma-tubulin, a component of the gamma-tubulin ring complex which is thought to be the main nucleator of microtubules in most cells [7]. Although gamma-tubulin is highly abundant in MTOCs in the apical domain of typical animal epithelia [51], to our surprise it was mostly undetectable in the apical domain of the distal tegument. Only small and sparse specks could be detected at a level around or above the microtriches (Fig 2D), which may actually represent the MTOCs of ciliated sensory receptors. In contrast, gamma-tubulin foci were common in the tegumental cytons (Fig 2E).

Altogether, the dearth of actin filaments in the distal tegument and the overall organization and abundance of microtubules suggest a possible role for the latter in polarized transport within this syncytium. However, the polarity of tegumental microtubules is not clear, and the mechanisms of organization and nucleation may be different from those found in typical epithelia.

### Protein synthesis is restricted to the tegumental cytons

Classic studies with radioactively labeled amino acids in other parasitic flatworms indicated that protein synthesis is largely absent from the distal tegument. However, some signal was always detected in the distal tegument in these studies, which may have been due to the inevitable diffusion of newly synthesized cytoplasmic proteins within the time frame of the labeling. Furthermore, the localization of protein synthesis has not been addressed in *M. corti* before. Here, we first confirmed that ribosomal RNA is restricted to the cytons and to the cytoplasmic bridges, by means of immunofluorescence and *in situ* hybridization (Fig 3A,B). Then, we localized the sites of new protein synthesis within the tegument using the ribopuromycylation method (RPM), in which we detected the incorporation of puromycin into nascent peptide chains in the ribosomes [52]. Puromycin was exclusively detected in the subtegumental region, confirming the absence of protein synthesis in the distal tegument (Fig 3C, S2). Altogether, these results implicate the existence of a transport mechanism for new proteins from the cytons to the distal tegument.

**Fig 3.**
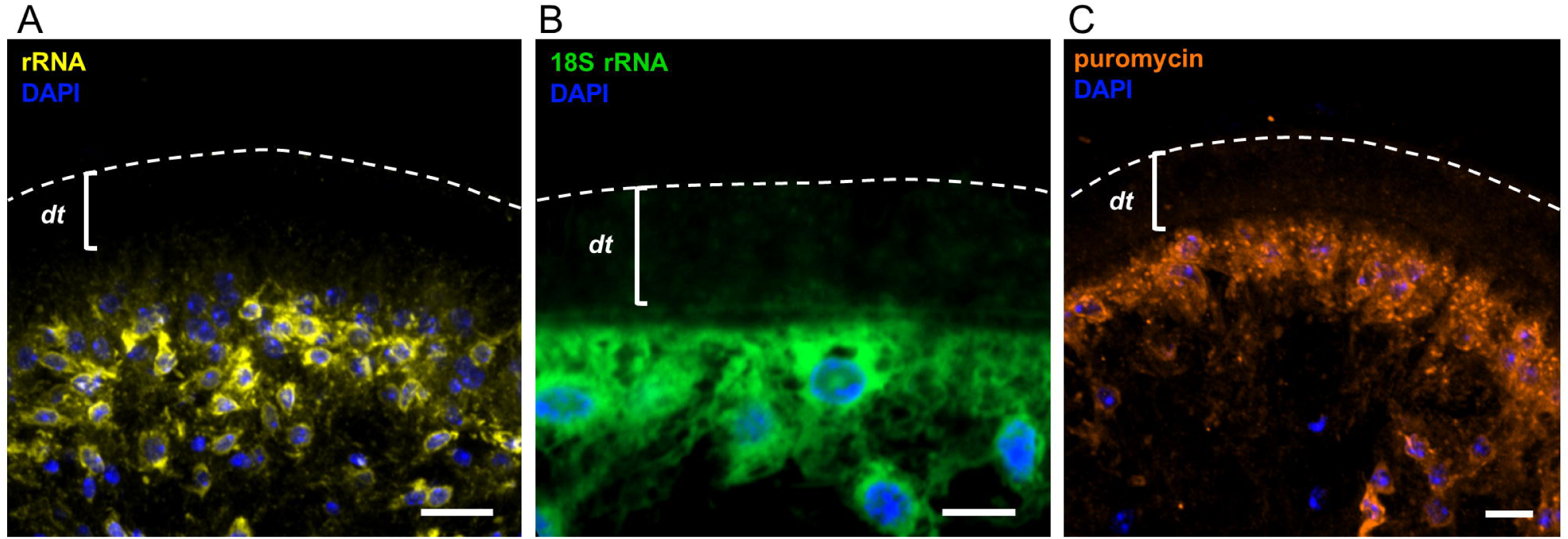
Protein synthesis in the tegument is restricted to cytons. (A) Immunofluorescent detection of rRNA. This experiment was performed twice. (B) 18S rRNA detection by *in situ* hybridization. This experiment was performed twice. (C) Immunofluorescent localization of puromycin incorporation (RPM). This experiment was repeated using two independent *in vitro* cultures of larvae with similar results. dt, distal tegument. Scale bars: 10 μm.

### Albendazole and albendazole sulfoxide eliminate microtubules in the distal tegument

We studied the effects of exposing *M. corti* larvae to 1 μM ABZ or ABZSO for 6 hours. This ABZSO concentration is similar to that found in the sera of patients treated for cestodiases, and the short time of exposure was selected in order to observe early effects of the drugs, which are more likely to be direct, rather than their long-term effects. Previous studies have shown that *M. corti* larvae are not overtly affected by concentrations of ABZ up to 20 μM for several days *in vitro* [53,54,55], and we determined that no loss of viability was caused under our *in vitro* conditions by 1 or 10 μM ABZ for at least three days (Fig S3).

Exposure to ABZ or ABZSO resulted in the complete loss of microtubules in the distal tegument (Fig 4A,B), while no effect was observed in solvent controls with dimethyl sulfoxide (DMSO). A decrease in the abundance of microtubules in comparison with the vehicle control was also apparent in the tegumental cytons (identified by dextran labeling) (Fig 4C). In contrast, microtubules were still abundant in the cytoplasm of other cells in the subtegumental region, and could also be detected in the sensory processes that traverse the distal tegument (Fig 4A-C). Increased concentrations and extended incubations with 10 μM ABZ for 18 hours also resulted in a strong depletion of microtubules in the tegument, but not in the nerve cords (Fig 4D). These results show that tegumental microtubules are strongly and specifically affected by low concentrations of ABZ and ABZSO. In comparison, we also assessed the effects of colchicine, a classic drug that potently and pseudo-irreversibly binds tubulin and inhibits microtubule polymerization in mammalian cells [56]. However, colchicine has been shown to have lower affinity for tubulin in diverse parasite species [57,58]. Treatment of *M. corti* larvae with 10 μM colchicine for six hours did not result in any discernible effects on tegumental microtubules, and the microtubules of the distal tegument only disappeared when larvae were treated with 100 μM colchicine (Fig 4A).

**Fig 4.**
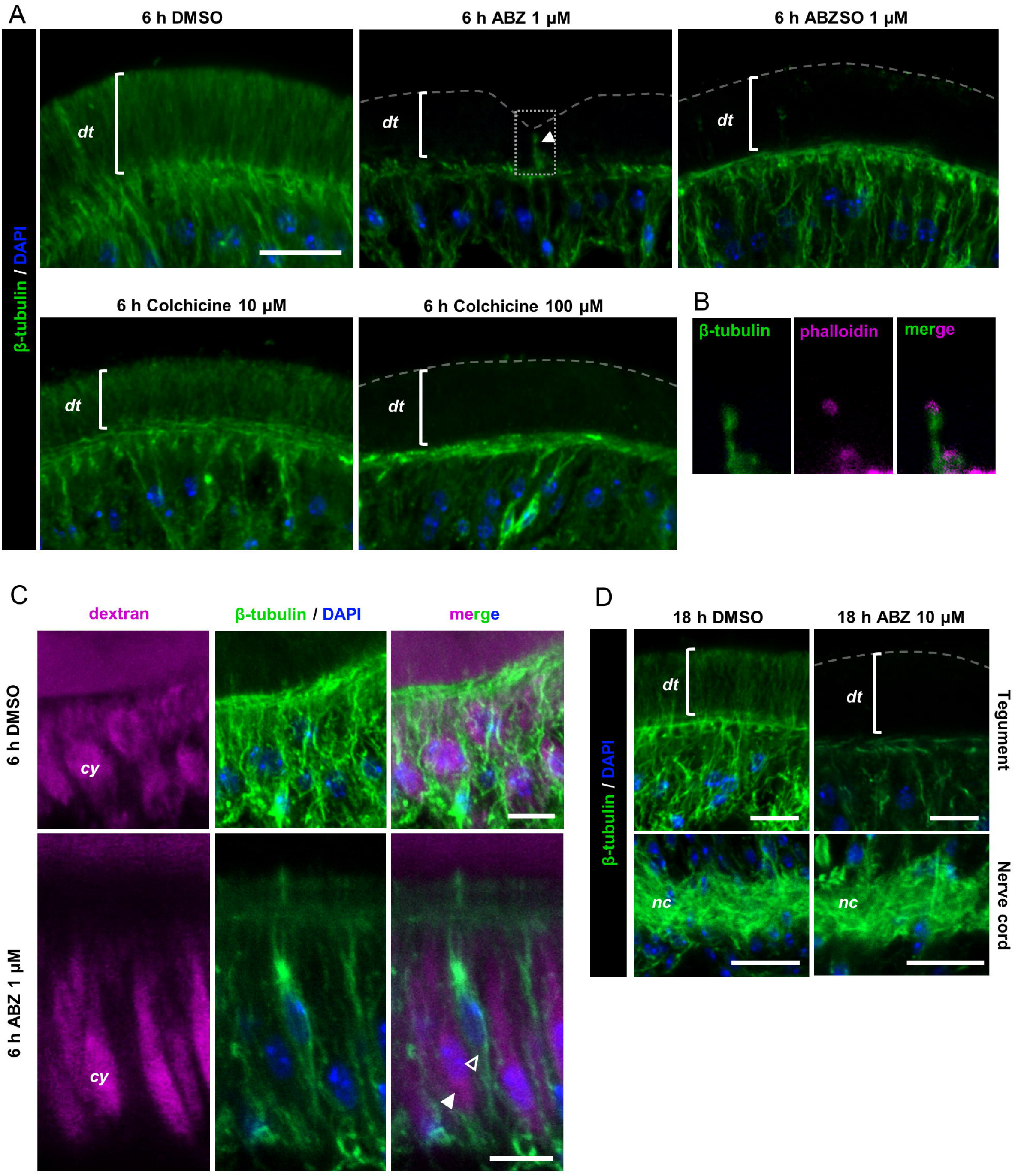
Effect of ABZ on tegumental microtubules. (A) Immunofluorescent detection of beta-tubulin after 6 h of *in vitro* treatment with ABZ or ABZSO (1 µM), colchicine (10-100 µM) and in DMSO controls. A sensory nerve ending traversing the tegument is indicated with an arrowhead in the upper middle panel. This experiment was repeated three times with different batches of larvae with similar results. (B) Inset of the dotted square in the upper middle panel of (A) showing the actin rings at the apical end of the nerve endings. (C) Immunofluorescent detection of beta-tubulin in the subtegumental region in control conditions (DMSO) and after ABZ treatment. Tegumental cytons are labeled with fluorescent dextran. A filled arrowhead indicates a tegumentary cyton, and the open arrowhead indicates a non-tegumental cell. This co-labeling experiment was performed once. (D) Immunofluorescent detection of beta-tubulin in the tegument (upper panels) and in the nerve cords (lower panels) after 18 h of *in vitro* incubation with 10 µM ABZ and in DMSO controls. This experiment was performed twice using different batches of larvae. Scale bars: A and D upper panels: 10 μm; C and D lower panels: 5 μm.

### Albendazole affects the incorporation of newly synthesized proteins in the distal tegument

Because of the extensive loss of microtubules in the tegument of ABZ treated larvae, it is possible that intracellular traffic could be affected. We determined the localization of secretory material within the tegument of treated and control parasites. To this end, we used the lectin Wheat Germ Agglutinin (WGA), which labels glycoconjugates that are found on the surface and distal tegument of *M. corti* larvae [59]. Treatment with 1 μM ABZ or ABZSO for 6 hours resulted in the accumulation of WGA^+^ material in the tegumental cytons, indicating that intracellular transport was affected (Fig 5A-C). In contrast, no effect was observed in the distribution of WGA staining after larvae were treated for 6 hours with 10 μM colchicine (Fig 5A-C).

**Fig 5.**
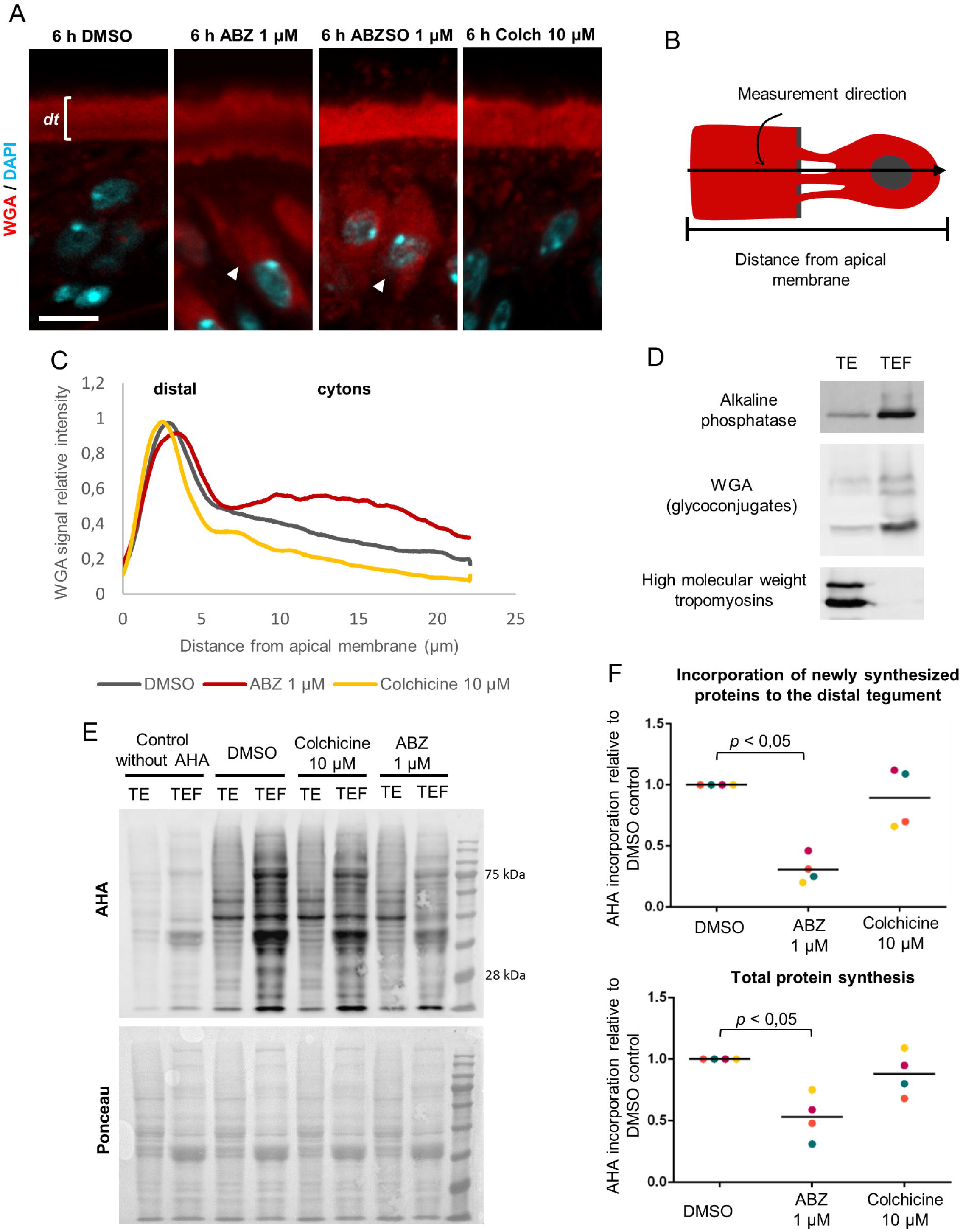
Effects of ABZ on the incorporation of new proteins into the distal tegument. (A) WGA labeling after incubation of larvae with 1 µM ABZ, 1 µM ABZSO, 10 µM colchicine and in DMSO controls. ABZ and ABZSO incubations resulted in increased WGA labeling in the tegumental cytons (arrowheads). dt, distal tegument. This experiment was repeated three times using different batches of larvae (B) Schematic drawing of the quantification of WGA staining in the tegument. Pixel intensity was measured along a line perpendicular to the apical region of the distal tegument. Each intensity value was normalized by dividing it by the maximum value of each condition. A total of 36 lines of 12 different parasites were measured and averaged. (C) Quantification of WGA labeling across the tegument after ABZ, colchicine and DMSO incubations. The positions of the distal tegument and the cytons are indicated. (D) Western Blot of tegumental enriched fractions (TEF) and total extracts (TE) for the detection of distal tegument markers (alkaline phosphatase and WGA+ glycoconjugates), and muscle fiber markers (high molecular weight tropomyosins). This experiment was repeated four times, starting from different batches of larvae, with similar results. (E) Detection of AHA labeling of proteins by Western Blot after incubation of larvae with ABZ 1 µM, colchicine 10 µM or DMSO controls (upper panel). The AHA signal in each lane was normalized using Ponceau S staining (lower panel). (F) Quantification of AHA labeling in TEFs (upper panel) and TEs (lower panel). Colors represent different independent replicates, using different batches of larvae (n=4). Mann-Whitney test. Scale bar: 5 μm.

The change observed in the distribution of WGA^+^ material after ABZ exposure is compatible with an inhibitory effect on vesicular traffic from the cytons to the distal tegument. To further explore the effect of ABZ on the incorporation of newly synthesized proteins into the distal tegument, we developed a method combining the labeling of protein synthesis in *M. corti* with the amino acid analog L-Azidohomoalanine (AHA) [60], and subcellular fractionation of the tegument to obtain protein fractions enriched for the distal tegument (Tegument Enriched Fractions, TEFs). First, we determined that AHA incorporation could be detected in new proteins in total extracts of *M. corti* by Western Blot. Signal above the background was detectable after one hour of incubation of the larvae with AHA *in vitro*, but robust signal required at least three hours of incubation. Second, we adapted to *M. corti* a protocol for obtaining TEFs [61], and validated these fractions by Western Blot by comparing them to total protein extracts. TEFs were highly enriched for glycosylated proteins found in the distal tegument that could be detected with the lectin WGA, and for the tegumental protein alkaline phosphatase (Fig 5D, S4). In contrast, TEFs were strongly depleted of muscular tropomyosins (Fig 5D). Because muscle fibers are located directly below the distal tegument (Fig S4) [62], these results are a strong validation that the enrichment procedure is specific for the distal tegument, and not for other cells or structures located in the subtegumental region.

AHA incorporation was readily detectable in TEF proteins of control larvae after three hours, demonstrating the incorporation of newly synthesized proteins in the distal tegument within this time frame, similarly to classic results with radioactive precursors in the tapeworm *Hymenolepis diminuta* [4,5]. In contrast, incorporation of new proteins in TEFs decreased on average to 31% of controls after 6 hours of incubation with 1 μM ABZ (Fig 5E,F). Surprisingly, total protein synthesis, as determined by AHA incorporation in total extracts, also decreased in ABZ treated larvae, although only to 53% on average (Fig 5E,F). Our results therefore demonstrate that ABZ can impair the incorporation of new proteins to the distal tegument, and part of this effect may be caused by the unexpected global reduction in protein synthesis caused by the drug. In contrast, exposure of worms to 10 μM colchicine had no effect on total protein synthesis or on the incorporation of new proteins to the distal tegument (Fig 5E,F).

### The effects of albendazole persist after drug removal

In many different animal cell models, microtubules are quickly regenerated after the removal of polymerization inhibitors [56]. We analyzed in *M. corti* the effect of ABZ removal for 12 hours, after 6 hours of exposure to ABZ 1 μM. Unexpectedly, microtubules were never regenerated in the distal tegument in four different experiments (Fig 6A). Similarly, although total protein synthesis sometimes recovered or even increased after ABZ removal, the incorporation of new proteins into the distal tegument did not recover to the levels found in control worms (Fig 6B). Thus, the effects of ABZ in the tegument outlast the exposure of larvae to the drug.

**Fig 6.**
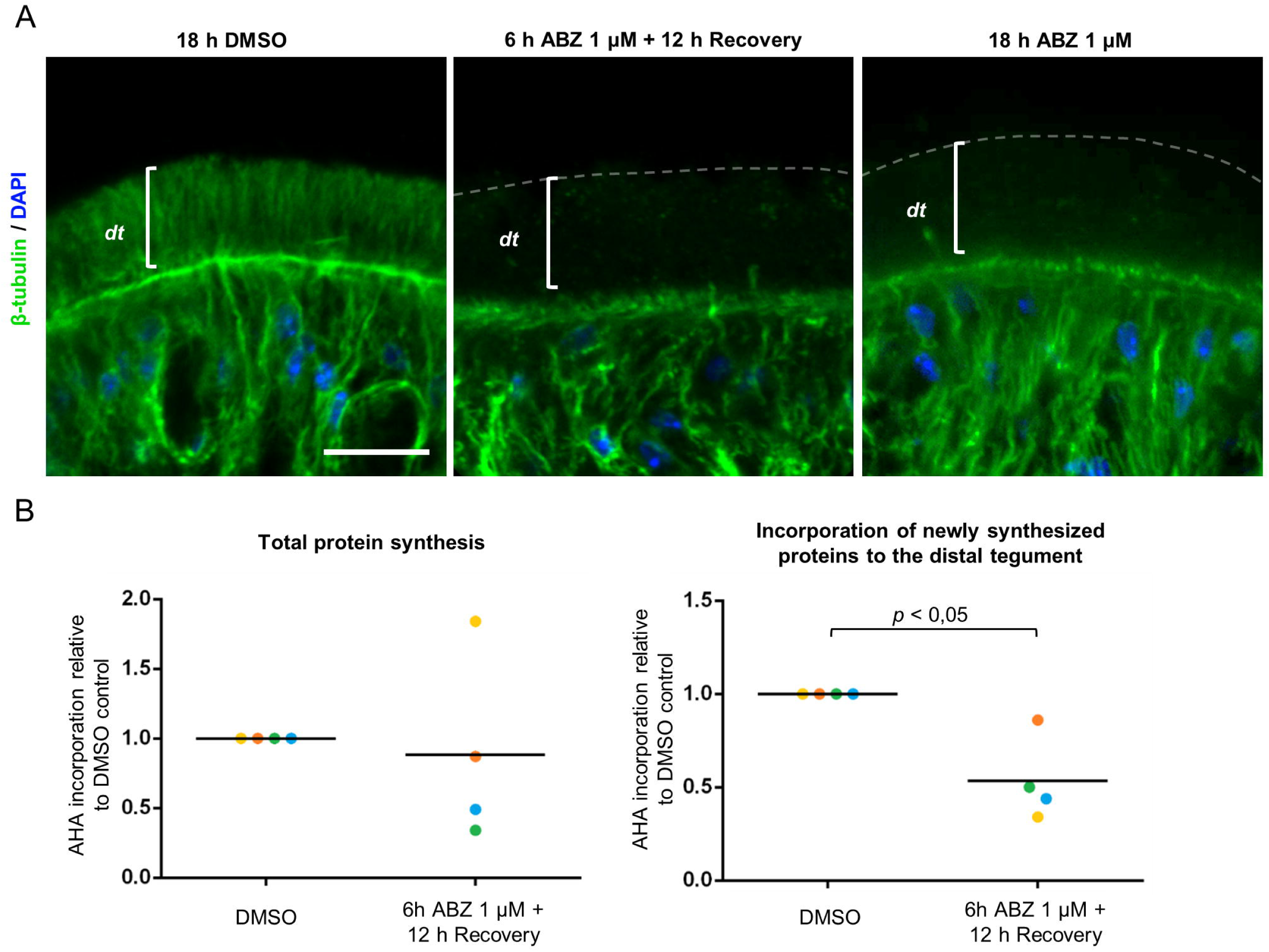
The effects of ABZ persist after drug removal. (A) Immunofluorescent detection of beta-tubulin after 6 h of ABZ incubation and 12 h of recovery (middle panel), 18 h of continuous ABZ incubation (right panel) and DMSO control (left panel). This experiment was repeated three times with similar results. (B) Incorporation of AHA in proteins in TEs (left panel) and TEFs (right panel) in DMSO controls, and after 6 h of incubation with ABZ 1 µM and 12 h of recovery. Values are from quantification by Western Blot as in Fig 5F. Colors represent independent replicates (n=4), using different batches of larvae. Mann-Whitney test. dt, distal tegument. Scale bars: 10 μm.

### Inhibition of protein synthesis is not correlated with activation of the unfolded protein response

The inhibition of protein synthesis caused by ABZ suggested that a general mechanism of translational inhibition could be activated by the drug. One highly conserved mechanism that inhibits translational initiation is the phosphorylation of the eukaryotic initiation factor eIF2α [63], which has been shown to occur in mammalian cell models after treatment with benzimidazoles [64,65]. Several pathways converge to eIF2α phosphorylation, including the Unfolded Protein Response (UPR), which is activated by unfolding and overcrowding of proteins in the endoplasmic reticulum [66], conditions that could be present in the tegumental cytons in which secretory traffic has been inhibited. Therefore, we analyzed the global levels of phosphorylated eIF2α (eIF2α-P) in larvae treated with ABZ. However, no increase in eIF2α-P levels was induced by 1 μM ABZ, and only a small increase was induced by 10 μM ABZ. In contrast, larvae treated with dithiothreitol (DTT), a known activator of the UPR, showed a 2.4-fold increase in the levels of eIF2α-P (Fig 7A).

**Fig 7.**
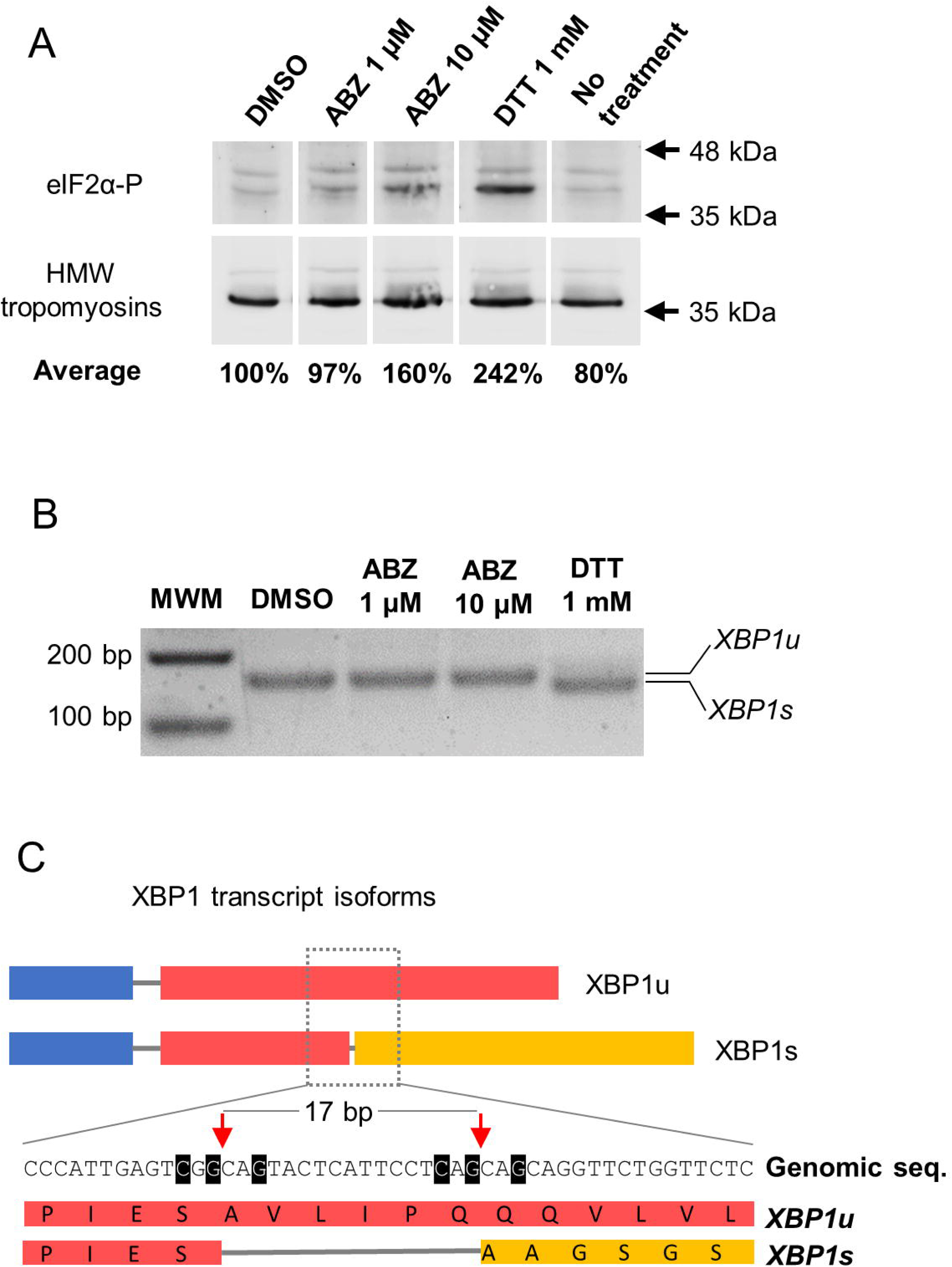
The UPR is not activated by ABZ. (A) Detection of phosphorylated eIF2α-P in total extracts by Western Blot. All lanes were cropped from the same blot. Quantification of eIF2α-P levels using high molecular weight (HMW) tropomyosins for normalization is shown for the average of three biological replicates (DMSO, ABZ 1 µM, ABZ 10 µM, no treatment) or two biological replicates (DTT). (B) RT-PCR of XBP1 transcripts. Only incubation with DTT resulted in the spliced form (XBP1s), which is 17 bp shorter than the unspliced isoform (XBP1u). This experiment was repeated three times with similar results, using different batches of larvae. (C) Schematic representation of XBP1 isoforms (not to scale) with an inset of the region surrounding the non-canonical intron, showing the 17 bp that are removed in *XBP1s*. Bases highlighted in black are identical to those essential for correct splicing in human XBP1. Red arrows indicate the splice sites, and orange background indicates the resulting frameshift.

In addition, we analyzed a different branch of the UPR, the activation of the transcription factor XBP1 by non-canonical alternative splicing of its mRNA mediated by IRE1/2 [66]. This splicing results in a longer open reading frame coding for an active protein. This pathway had not been demonstrated in flatworms so far. We identified XBP1 and IRE1/2 homologs in *M. corti* (MCU_007529 and MCU_001864, respectively), and showed that the classical UPR activator DTT induced non-canonical splicing in *M. corti* larvae that resulted in the removal of 17 bases in the *XBP1* mRNA (Fig 7B,C). Nucleotide positions that are essential for non-canonical splicing in humans [67] were conserved in *M. corti*, and the nucleotide sequence of the non-canonical intron was highly conserved in XBP1 homologs from the parasitic flatworms *Echinococcus multilocularis* and *Schistosoma mansoni*, as was the amino acid sequence of the altered open reading frame (Fig S5). However, ABZ at 1 or 10 μM did not induce non-canonical splicing of XBP1 (Fig 7B, C). Altogether, our results indicate that inhibition of protein synthesis by ABZ occurs independently of activation of the UPR.

## Discussion

### Organization of the cytoskeleton in the tegument

The tubulin cytoskeleton shows a regionalized organization in the syncytial tegument of *M. corti*. The abundance of microtubules in the cytoplasmic bridges that connect the cytons to the distal tegument was previously noted by electron microscopy in *M. corti* and in other parasitic flatworms [45,10], and suggests a role of microtubules in intracellular traffic between these subcellular domains. The abundance and apicobasal orientation of microtubules in the distal tegument is reminiscent of the polarized microtubule bundles found in animal epithelia [7], suggesting roles in polarized traffic, but their arrangement as radiating bundles originating from the cytoplasmic bridges is unique. On the other hand, the virtual lack of actin filaments in the distal tegument was also previously described in the cestode *Diphyllobothrium dendriticum* [68], and suggests that the actin cytoskeleton may not be important for intracellular trafficking in this region.

The polarity of microtubules in the distal tegument cannot be determined from our results, but the lack of gamma-tubulin in the apical domain suggests that this does not function as a MTOC in the tegument, opening the possibility that the minus end could be located at the basal region. In the trematode *Fasciola hepatica*, microtubules in the tegument have also been described by immunofluorescence, and in this case their abundance was low in the distal tegument, and they did not reach the apical region, which would be concordant with this possibility [69]. The bouquet configuration of distal microtubules is suggestive of microtubule branching. The best characterized mechanisms of microtubule branching in other animals are based on the recruitment of gamma-tubulin to pre-existing microtubules [70,7], but there is no gamma-tubulin at the possible branching sites in the basal region of the distal tegument of *M. corti*. However, gamma-tubulin independent mechanisms of nucleation and branching have been recently proposed in other models [71,72].

### Tegumental microtubules as targets of benzimidazoles

The microtubules of the tegument were exquisitely sensitive to ABZ and ABZSO, whereas little to no effect was apparent in other cells. This suggests that the tegument of *M. corti*, and perhaps of other cestodes, is one of the main tissular targets of benzimidazoles. The effect of ABZ on the tegument of *M. corti* was long-lasting even after removal of the drug. This could be due in part to the high stability of binding between benzimidazoles and tubulin in helminths [13], and may be relevant for chemotherapy, since drug concentrations in plasma decrease notably between successive doses [34]. In contrast, colchicine showed very low potency against tegumental microtubules, which is concordant with the low affinity of this drug for tubulin in different invertebrate species [57], and with the lack of efficacy in a rodent model of infection by *E. multilocularis* [73].

There are several possible explanations for the differential sensitivity of the tegument to benzimidazoles. First, the tegument is the tissue that is most exposed to drugs in the external media. However, the microtubules of sensory receptors, which are similarly exposed to the external medium, did not appear to be affected by ABZ or ABZSO. A second, perhaps more likely explanation could be related to differential dynamics of microtubules in different tissues. For example, microtubules in the sensory receptors are likely to be highly stable, as suggested by the abundance of acetylated tubulin. In contrast, microtubules in the distal tegument are likely to be highly dynamic, which would render them more susceptible to the effects of a drug inhibiting tubulin polymerization. Finally, there are several different isoforms of beta-tubulin in parasitic flatworms, and some of these have amino acid substitutions that are associated with benzimidazole resistance in other helminths. In *Echinococcus multilocularis*, proliferating stem cells are thought to express a resistant beta-tubulin isoform, which could explain the lack of efficacy of ABZ against this crucial cell population [74]. It is possible that differential expression of tubulin isoforms could also contribute to the preferential effects of ABZ and ABZSO in the tegument of *M. corti*.

Interestingly, ABZ and ABZSO had similar effects at low concentrations in the tegument of *M. corti*, and both compounds have similar potency *in vitro* against *Echinococcus* spp. larvae [27,28]. This contrasts with reports in mammalian cells *in vitro* and in other vertebrate models, for which ABZ is invariably much more toxic than ABZSO [75,76,77,78]. Because ABZSO is the main active metabolite found in the plasma of patients, this could be one more element participating in the selectivity of ABZ treatment against cestode parasites. In any case, benzimidazole chemotherapy against *Echinococcus* spp. requires long treatments and is only parasitostatic [22,79]. The short-term effects *in vitro* are subtle at low concentrations, and it is possible that the effects of benzimidazole treatment on tegument maintenance and turnover *in vivo* could be synergistic with the damage caused on the parasite surface by the immune system. Indeed, there is evidence that benzimidazole treatment is ineffective in the absence of a functional adaptive immune system [80].

### Effects of ABZ on protein synthesis and tegumental turnover

The effect of ABZ on tegumental microtubules correlated with the accumulation of secretory material in the cytons, and with the inhibition of the incorporation of new proteins in the distal tegument. Our results suggest that disrupting the microtubule cytoskeleton affects intracellular trafficking of new proteins from the cytons to the distal tegument. However, we also observed an unexpected effect of ABZ on total protein synthesis, which does not allow us to dissect how much of the inhibition in the incorporation of new proteins in the distal tegument is due to disrupted traffic, versus how much is an effect of inhibited protein synthesis. An effect on protein synthesis and secretion has sometimes been observed after microtubule depolymerization in mammalian cell models [9], including recent studies showing the activation of eIF2α phosphorylation after benzimidazole treatment [64,65]. In contrast, in *M. corti* ABZ did not activate the UPR and the inhibition of protein synthesis appears to be independent of eIF2α phosphorylation. To the best of our knowledge, inhibition of protein synthesis by benzimidazoles in helminths has only been shown for triclabendazole sulfoxide (TCBZSO) in *F. hepatica* [81]. Importantly, it is thought that TCBZSO may be one of the few benzimidazoles that do not target microtubules, as it does not bind *F. hepatica* tubulin *in vitro* [82], and resistance to the parent drug triclabendazole in *F. hepatica* is not genetically linked with beta-tubulin isoforms [83]. Future work with different benzimidazoles in other helminth species will be important to determine if the effect of inhibition on protein synthesis may be a general effect of this class of drugs.

## Materials and Methods

### Animals

*M. corti* (syn. *vogae*) tetrathyridia larvae from the strain originally isolated by Specht and Vogue [84] were obtained from peritoneal infection of C57BL/6 mice and Sprague Dawly rats, in collaboration with the Laboratorio de Experimentación Animal, Facultad de Química, Universidad de la República, Uruguay (“Mantenimiento del cestodo *Mesocestoides vogae* para estudios de actividad antihelmíntica y uso de otros servicios de Udelar”; protocol number 10190000011119, approved by Comisión Honoraria de Experimentación Animal, Uruguay). The parasites were collected at three to six months post-infection.

For immunofluorescence experiments where a mouse primary antibody was used, larvae obtained from rats were utilized in order to use an anti-mouse secondary antibody, avoiding nonspecific signal from host antibodies adhered to the larvae surface. When rabbit or rat primary antibodies were used, larvae obtained from mice were utilized.

### In vitro culture

Tetrathyridia larvae were obtained under sterile conditions from the peritoneum of mice or rats and kept at 4°C on sterile Phosphate Buffered Saline (PBS) until use for up to 7 days. Before conducting any experiments the parasites were cultured *in vitro* overnight (at least 12 h) in RPMI-1640 medium (Sigma, R4130, United Kingdom) containing antibiotic-antimycotic (Thermo, 15240-062, U.S.A.), supplemented with 10% fetal bovine serum (FBS; Capricorn, cat N° FBS-11A, collected in South America), at 37°C and 5% CO2.

### Fixation and cryosections

Tetrathyridia were washed with PBS to remove traces of medium and fixed overnight at 4°C with 4% paraformaldehyde (PFA) dissolved in PBS, with gentle shaking. After extensive washing with PBS, the larvae were then embedded in Tissue-Tek OCT compound (Sakura FineTek cat. no. 4583, U.S.A.), immediately frozen with liquid nitrogen, and used to cut 10 μm cryosections.

### Dextran labeling

For fluorescent labeling of the tegument, we adapted the protocol published by Wendt et al [47] for the trematode *Schistosoma mansoni*, using a fluorescent fixable dextran (Thermo, D3312, U.S.A.). Larvae were cultured *in vitro* overnight. Afterwards, worms were transferred to a 1.5 ml centrifuge tube wrapped in aluminium foil, and vortexed for three minutes in a solution of 2 mg/mL of dextran dissolved in distilled water (at least 5 volumes of labeling solution per volume of larvae). The stained larvae were quickly washed three times with PBS. At this point, tetrathyridia were either fixed or put back in culture overnight in the same conditions indicated before, and fixed afterwards. Immediate fixation after shock produced uneven labeling of the tegument. To improve homogeneity of the label, and to allow the tegument to recover, we added an overnight culture step after the shock, which gave a homogeneous signal. This method was used for experiments combining dextran labeling and immunofluorescence. Larvae were stained with 4′,6-diamidino-2-phenylindole (DAPI; Sigma, D9542, Israel) and analyzed by confocal microscopy in whole-mounts, or included in OCT for posterior cryosectioning and combination with immunofluorescence experiments.

### Immunofluorescence

Experiments were conducted on cryosections of *M. corti* adapting the protocol described in Koziol et al [38]. Cyosections were washed with PBS with 0,1% Triton (PBS-T) and blocked with PBS-T plus 1% Bovine Serum Albumin (BSA; Capricorn, U.S.A.) and 5% sheep serum (Sigma, S3772, U.S.A.). Incubations with the primary and secondary antibodies were done for 72 h each, in PBS-T with 1% BSA and 0,02% sodium azide. This extended incubation was necessary to ensure the adequate penetration of the antibodies in the distal tegument. After incubation with each antibody, cryosections were extensively washed with PBS-T. Staining with DAPI and phalloidin-iFluor 488 (Abcam, ab176753, U.S.A.) was performed together with the secondary antibody incubation. The specimens were mounted with Fluoroshield (Sigma, F6182, U.S.A.). The antibodies used and their dilutions are provided in Table S1. Tubulin antibodies were validated by Western Blot (Fig S6). All immunofluorescence experiments were repeated at least twice.

### Alkaline Phosphatase histochemistry

Alkaline phosphatase activity was detected in cryosections with nitro blue tetrazolium chloride and 5-bromo-4-chloro-3-indolyl phosphate (Sigma, 72091, Germany) as described by Cox and Singer [85]. Slides were mounted using 80% glycerol with Tris-HCl 50 mM, pH 8.0.

### Detection of glycoconjugates with WGA

Cryosections were washed with PBS and incubated in the dark for 72 h with PBS-T plus 1% BSA, 10 μg/mL fluorescent Wheat Germ Agglutinin (WGA, Vector Labs, U.S.A.), DAPI and 0,02% sodium azide. The slides were then washed with PBS-T and mounted with Fluoroshield medium (Sigma, F6182, U.S.A.).

### Whole Mount *In Situ* Hybridization

*M. corti* gDNA was used to amplify a fragment of the 18S rRNA gene by PCR. Primers were slightly modified from those published by Muehlenbachs et al 2015: 5’-GGGGATGGGTGCACTTATTAGA - 3’, 5’ - GTTATCACCATGGTAGGCAGGT - 3’. The amplicon was cloned into pCR II vector (Dual promoter Kit, Invitrogen, 45-0007LT, U.S.A.) and digoxigenin-labeled RNA probes were generated by *in vitro* transcription using SP6 or T7 polymerases (Thermo, EP0131 and EP0111 respectively, Lithuania) in reactions containing 3,5 mM digoxigenin-UTP, 6,5 mM UTP, and 10 mM CTP, ATP and GTP.

Fixation and whole mount in situ hybridization were performed as described by Koziol et al [86], except that permeabilization was done with 30 μg/mL Proteinase K (New England Biolabs, P8102S, U.S.A.) for 40 minutes.

### RPM

We adapted the protocol published by David et al [52]. *M. corti* larvae were incubated in RPMI medium supplemented with 10% FBS plus 92 μM puromycin (Millipore, 540411, China), at 37°C, for 30 minutes. As a control, larvae were pre-incubated in the presence of 9,4 μM anisomycin (Sigma, A9789, Germany) for 15 minutes before puromycin was added to the medium. After incubation, the larvae were fixed and processed for cryosections after which puromycin detection was performed by immunofluorescence, as indicated before. We also performed experiments of co-incubation with puromycin and emetine (Millipore, 324693, China) as described originally by David et al [52], in order to prevent the release and diffusion of truncated puromycylated peptide chains, with similar results (Fig S2).

### Drug treatments

Incubations of *M. corti* larvae with different drugs were performed in RPMI medium supplemented with 10% FBS at 37°C for 6 h. The drugs used were albendazole (ABZ, Sigma A4673, China), albendazole sulfoxide (ABZSO; ChemCruz, sc-205838, U.S.A.) and colchicine (Sigma, C9754, India), which were compared with incubations with 0.01% DMSO (Sigma, D-8779, U.S.A.) as vehicle control. Longer incubations for 18 h in the presence of 10 μM ABZ were compared with 0.1% DMSO as vehicle control. For the evaluation of recovery after drug removal, larvae were incubated with the drug for 6 h and washed extensively in culture media, followed by 12 h of recovery incubation in culture medium without the drug.

### Viability assay

Parasites were cultured in RPMI medium supplemented with 10% FBS at 37°C and 5% CO2 for 10 days in the following conditions: 1 μM ABZ, 10 μM ABZ and 0,1% DMSO. Viability was evaluated by eosin staining as described by Fabbri et al [87]. Non-viable worms were defined as those that failed to exclude the dye, independently of their motility.

### Isolation of Tegument Enriched Fractions and total protein extracts

We adapted to *M. corti* larvae protocols that were previously published for other cestode species [88,61]. After washing with PBS, one volume of larvae was incubated with 3 volumes of PBS plus 0.2% Triton X-100 (Baker, U.S.A.) on ice and with mild shaking. Immediately after, larvae were vortexed for 30 seconds to mechanically detach the distal tegument. The “naked” worms were pelleted by centrifugation at 200 g, then the supernatant was recovered and centrifuged at 2500 g. The pellet obtained after this centrifugation step is the Tegument Enriched Fraction (TEF). We washed the pellet with PBS, re-centrifuged the material at 2500 g, discarded the supernatant and stored the dry pellet at -20°C until use. All centrifugation steps were done in a refrigerated centrifuge, at 4°C.

For protein quantification TEFs were resuspended in incomplete Loading Buffer (without β-mercaptoethanol and Bromophenol blue) plus protease inhibitor cocktail (Sigma, P8849, Germany), boiled at 100°C for 10 minutes and quantified using Pierce™ BCA Protein Assay Kit (Thermo, 23227, U.S.A.).

To obtain total protein extracts, larvae were snap frozen in liquid nitrogen until further use. For pestle homogenization, 2 volumes of incomplete Loading Buffer with protease inhibitor cocktail were added to 1 volume of worms. Homogenization was conducted on ice, after which the homogenate was boiled at 100°C for 10 minutes and centrifuged for 5 minutes at 17.000 g. Supernatant was recovered and quantified as indicated for TEFs.

### Western Blot

After adding β-mercaptoethanol and bromophenol blue, each sample (20 μg) was run in a 10% SDS-PAGE gel. Proteins were then transferred to a nitrocellulose membrane (Thermo, 88018, Germany). The membrane was blocked with either 5% BSA in Tris-NaCl plus 0,1% Tween (Sigma, P9416, U.S.A.) (TBS-T) for detection of phosphorylated eIF2α, or 5% milk in TBS-T for all other proteins.

After blocking, an overnight incubation with the primary antibody (Table S1) in blocking solution was performed at 4°C with mild agitation. Then, after washing with TBS-T, incubation with the secondary antibody conjugated to Horseradish Peroxidase (HRP) in blocking solution was conducted at room temperature, for 1 h with mild agitation. Finally, the membrane was washed with TBS-T and developed using the chemiluminescent reagent SuperSignal™ West Pico PLUS (Thermo, 34580, U.S.A.). Imaging was done again with a G:BOX system.

For the detection of glycoconjugates, the membrane was incubated with lectin WGA conjugated to rhodamine (Vector Labs, U.S.A.) in TBS-T for 1 h at room temperature, with mild shaking, in the dark. Imaging of fluorescence was done with the G:BOX system.

### AHA labeling and quantification

*M. corti* larvae were cultured in RPMI medium supplemented with 10% FBS at 37°C for 2.5 h with 1 μM ABZ, 10 μM colchicine or 0.01% DMSO, after which they were pre-incubated in RPMI medium without methionine (-Met) nor FBS (-FBS) (Gibco, A14517-01, U.S.A), plus the drug for 30 min. Then, the medium was replaced with RPMI (-Met -FBS) with the drug plus 50 μM of the methionine analog L-Azidohomoalanine (AHA, Sigma, 900892, U.S.A.) and parasites were incubated for 3 h at 37°C. Control larvae were cultured for 6 h in the same media without any drug or AHA.

After AHA labeling, larvae were processed to obtain total protein extracts and tegument enriched fractions, which were quantified, as indicated above.

AHA was detected by means of a Click-It reaction (Thermo, C10276, U.S.A.) using alkyne-Biotin (Sigma, 764213, U.S.A.), following the instructions of the manufacturer. The biotin labeled proteins were re-dissolved with incomplete loading buffer with extensive vortexing and heating to 70 °C, and the concentration of the labeled proteins was determined as described above.

Afterwards, the proteins were run in a 10% SDS–PAGE gel and transferred to nitrocellulose membranes (Thermo, 88018, Germany), and total proteins were stained with Ponceau S and imaged. The membrane was blocked with 5% non-fat milk in TBS-T, and the biotin label (corresponding to newly synthesized proteins) was then detected by incubating the blocked membrane with streptavidin-HRP (Thermo, S911, U.S.A.) in blocking solution overnight at 4°C with mild shaking, and developing with SuperSignal™ West Pico PLUS (Thermo, 34580, U.S.A.).

Quantification was conducted using the software Fiji [89]. A Region of Interest (ROI) was defined to contain each individual lane, then the Integrated Density was measured as the sum of the values of the pixels in the selected ROI. For the Ponceau S image, this represents the signal from total protein loaded in the lane, and for the AHA detection image, this represents the amount of newly synthesized proteins. The signal from a blank lane without any proteins (“PONCEAU blank”) was substracted from the signal from each experimental lane for Ponceau S staining (“PONCEAU drug”). The signal of a control lane containing proteins from worms not incubated with AHA (“No AHA ctrl”) was subtracted from the signal from each lane for AHA detection (“AHA drug”). The amount of specific AHA incorporation was defined as the ratio between the corrected AHA and Ponceau S signals. Then, relative quantification was performed, for both total protein extracts and TEFs, by comparing the results from drug treated parasites to DMSO control parasites (“AHA DMSO”, “PONCEAU DMSO”), according to the following equation: [(AHA drug - No AHA ctrl) / (PONCEAU drug - PONCEAU blank)] / [ (AHA DMSO - No AHA ctrl) / (PONCEAU DMSO - PONCEAU blank)].

### Analysis of XBP1 splicing

We identified *M. corti* XBP1 and IRE1/2 orthologs by BLAST using the human genes as queries (NCBI accession numbers NP_005071.2 and NP_001424.3, respectively), against the predicted proteome of *M. corti* (version WBP14), which was downloaded from WormBase Parasite [90]. RNA was extracted from *M. corti* larvae after incubations with 1 μM ABZ, 10 μM ABZ and 0,1% DMSO for 6 h, and after incubation with 1 mM DTT for 2 h (longer treatments killed the parasites), using the Zymo DirectZol RNA Miniprep Kit (R2070, U.S.A.). RNA (500 ng) from each experimental condition was reverse transcribed with SuperScript II (Thermo, 100004925, U.S.A.), and the resulting cDNA was used as template for amplification by RT-PCR with HighTaq (Bioron, 111105, Germany), using the following primers: 5’-GAGGAGACAGATTGAGCCCATTG-3’, 5’-ATCAACATTCCCACTACAATTAAGCG-3’.

This reaction amplified a 144 bp fragment of the *M. corti* XBP1u cDNA surrounding the non-canonical splice site, which was identified based on conserved nucleotides that are essential for its splicing in mammals, and on the position of stop codons in the original reading frame. The current automatically generated gene model for MCU_007529 has a canonical intron incorrectly predicted in this region, which was rejected by our RT-PCR data. Amplicons were analyzed by electrophoresis in 2% agarose gels. Gel purification and Sanger sequencing (Macrogen, Korea) was performed for an amplicon from each condition. Secondary structure prediction of the region of the non-canonical intron was performed based on alignments of the sequence of *M. corti XBP1* together with the equivalent regions from *E. multilocularis XBP1* (EmuJ_000843400) and *S. mansoni XBP1* (Smp_335350) with LocARNA-P [91].

### Microscopy

Samples were imaged by confocal microscopy (Zeiss LSM 800CyAn and LSM 980 with an Airyscan detector, Advanced Bioimaging Unit of the Institute Pasteur of Montevideo).

### Statistics

Mann-Whitney test (two-tailed) was carried out using Past4.0 software [92] (significance was considered at p < 0.05). For this non-parametric test, a sample size of four replicates was chosen to detect significant differences if the effect size was large enough and the variance small enough to prevent overlap between samples. No test for outliers was performed, and no data points were excluded in any experiments. No exclusion criteria were predetermined for any samples and larvae were arbitrarily selected for the different experimental treatments. GraphPad Prism software, v. 8.0.2 (Graphpad Software, Inc., San Diego, CA, USA) was used for plots.

## Acknowledgements

The authors thank Beatriz Munguía, Laboratorio de Experimentación Animal, Facultad de Química, Universidad de la República, Uruguay for providing *M. corti* larvae. The authors gratefully acknowledge the Advanced Bioimaging Unit at the Institut Pasteur Montevideo for their support and assistance in the present work. The authors would also like to acknowledge the help of Rosario Rovira in the making of the wind rose histogram.

## Data and resource availability

All relevant data and resources can be found within the article and its supplementary information.

## Supporting information

**S1 Fig. Organization of the tegument of *M. corti*.** (A) Co-labeling of the tegument and muscle fibers with dextran and phalloidin, respectively, showing how cytoplasmic bridges traverse the outer muscle layer in between the muscle fibers, just below the distal tegument. Not all cells in the subtegumental region are tegumental cytons: the filled arrowhead indicates a dextran^+^ cyton, and the open arrowhead indicates a dextran^-^ nucleus. (B) We found two distinctive morphologies of putative sensory nerve endings, both of which appear to end in a cilium-like protrusion. One sensory receptor has a dilated apical region (shown in Fig 2B in the main text) while the other, shown here by immunofluorescence for acetylated tubulin, does not have a dilation and appears to be surrounded by a depressed conical region of the distal tegument. (C) Schematic drawings of the two sensory nerve ending morphologies traversing the tegument. The possible position of septate junctions, based on phalloidin staining and previous electron microscopy descriptions are represented in purple, with black lines. dt, distal tegument; br, cytoplasmic bridges; cy, cytons; dt, distal tegument. Scale bars: A: 10 μm; B: 5 μm.

**S2 Fig. Controls of the ribopuromycylation experiment.** Emetine prevents the release of the puromycin labeled peptide from the ribosome, ensuring the observation of the protein synthesis sites (left panel). We obtained essentially identical results with and without emetine. A specificity control with anisomycin (which inhibits protein translation; middle panel) is shown, as well a control without puromycin (right panel). Scale bars: 10 μm.

**S3 Fig. Effects of ABZ on the viability of *M. corti*.** (A) Parasites were cultured *in vitro* in the presence of ABZ 1 µM or 10 µM. Viability was assessed by the eosin exclusion method. Viability was not significantly affected for at least 3 days, and then decreased to 38% and 10% for ABZ 1 µM and 10 µM, respectively, by day 10. (B) Detailed pictures of live parasites show blebs evidencing damage in the tegument of ABZ treated parasites on day 7. Scale bar: 200 μm.

**S4 Fig. Localization of tegument and muscle markers of *M. corti*.** (A) Localization of the enzyme alkaline phosphatase revealed by histochemistry on cryosections shows that it is mainly located in the distal tegument. (B) Detection of high molecular weight (HMW) tropomyosins show their location immediately below the distal tegument. dt, distal tegument. Scale bars: upper A: 100 μm; lower A and B: 10 μm.

**S5 Fig. Conservation of XBP1 in parasitic flatworms.** (A) Alignment of the region of XBP1 transcripts of *Echinococcus multilocularis* (Emu), *Schistosoma mansoni* (Sman) and *M. corti* (Mc) surrounding the 17 bp non-canonical intron (+/- 20 bp), labeled according to sequence conservation. The corresponding sequence of *Homo sapiens* XBP-1 is shown for comparison, including those residues that are essential for non-canonical splicing. (A) Structural prediction of the region of XBP1 transcripts from parasitic flatworms surrounding the non-canonical intron. The alignment in the left shows the conservation of the predicted secondary structure, which is shown on the right. The secondary structure consists of a double stem-loop structure similar to the unconventional splice junction of other species. (C) Protein sequence alignment of the spliced form of XBP1 (XBP1s) from Mc, Emu and Sman showing conservation at the amino acid level. Black dotted line marks the position of the frameshift produced by non-canonical splicing.

**S6 Fig. Western Blot of tubulin antibodies.** (A) Detection of alpha-tubulin shows a unique band of appropriate size in both soluble tegument fraction and total extracts of *M. corti.* (B) Detection of beta-tubulin shows two main bands in the middle range, the upper one has the expected size and the band immediately below could correspond to partial degradation of beta-tubulin in the extracts. (C) Detection of gamma-tubulin in total extracts of *M. corti* resulted in a single band of appropriate size. Total extract of HeLa cells was used as a positive control.

**S1 Table. Antibodies and their dilutions.**

