## Supplementary Material for "Albendazole specifically disrupts microtubules and protein turnover in the tegument of the cestode *Mesocestoides corti*"

Supplementary Figure 1

A

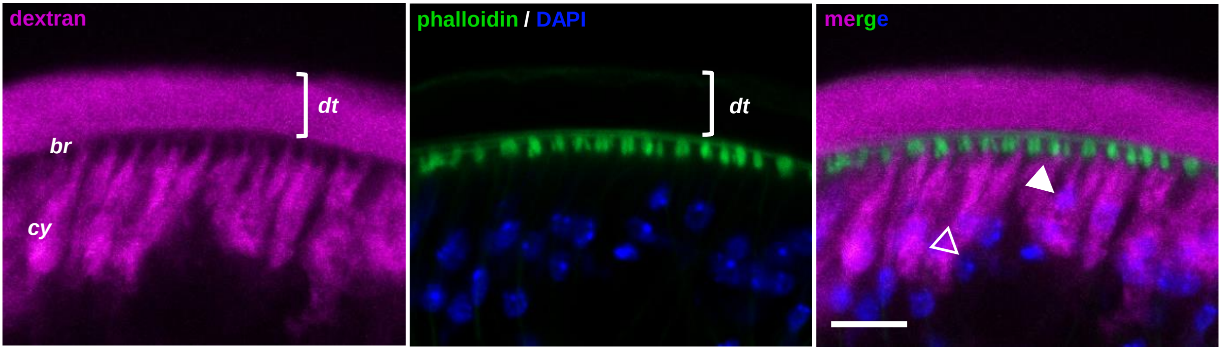

B

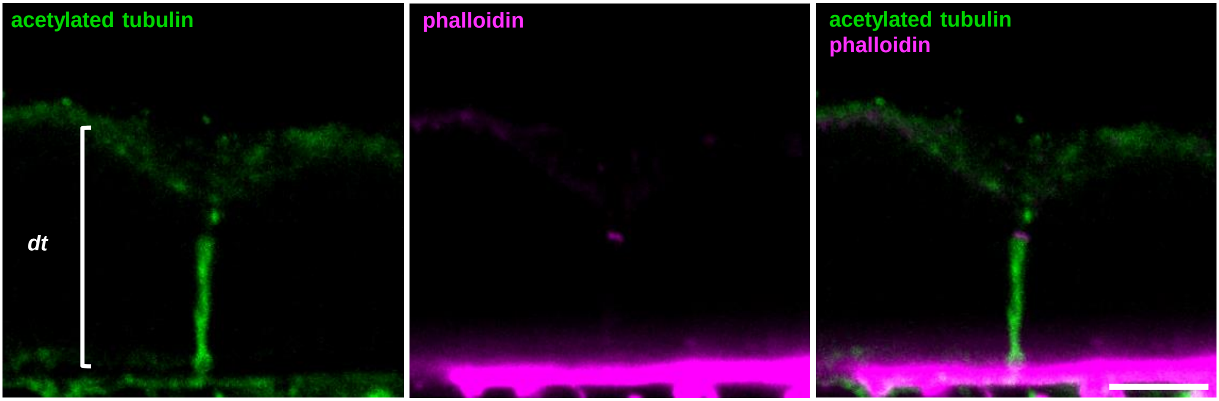

C

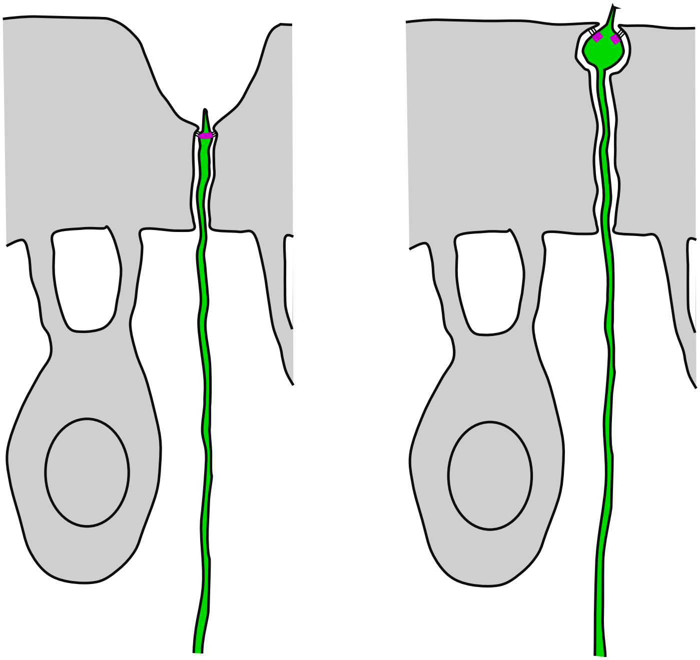

Supplementary Figure 2

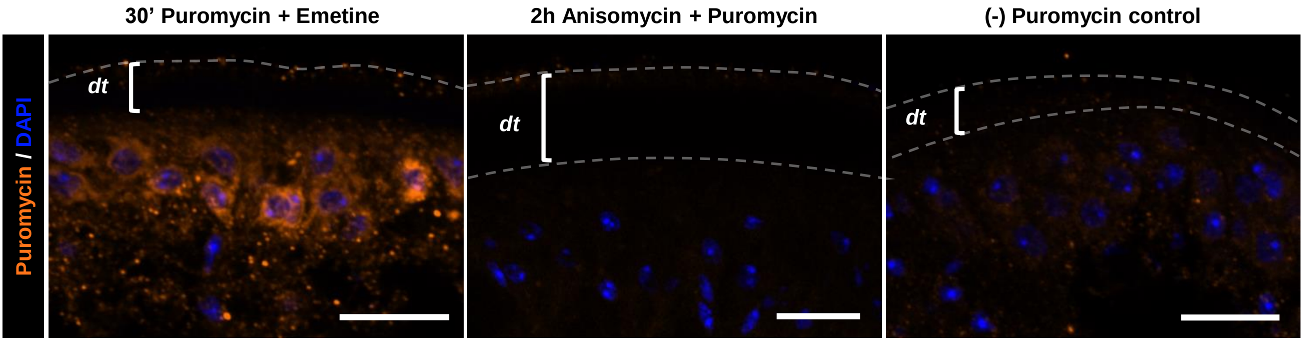

Supplementary Figure 3

A

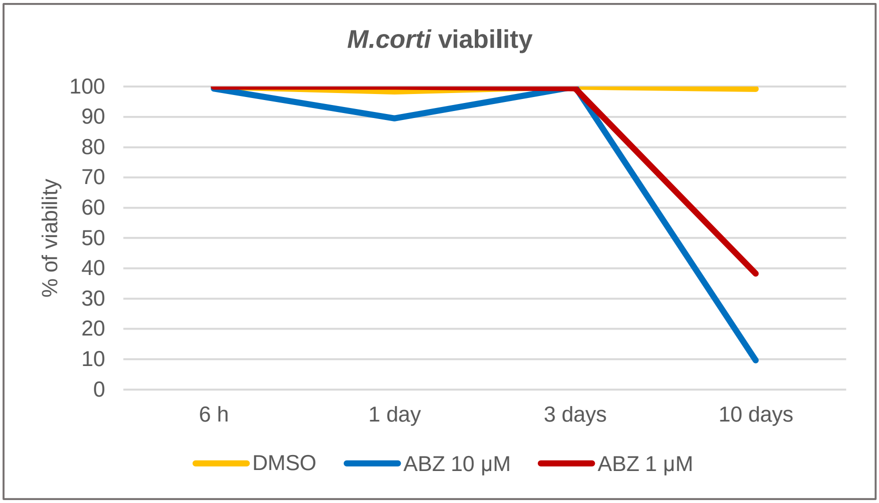

B

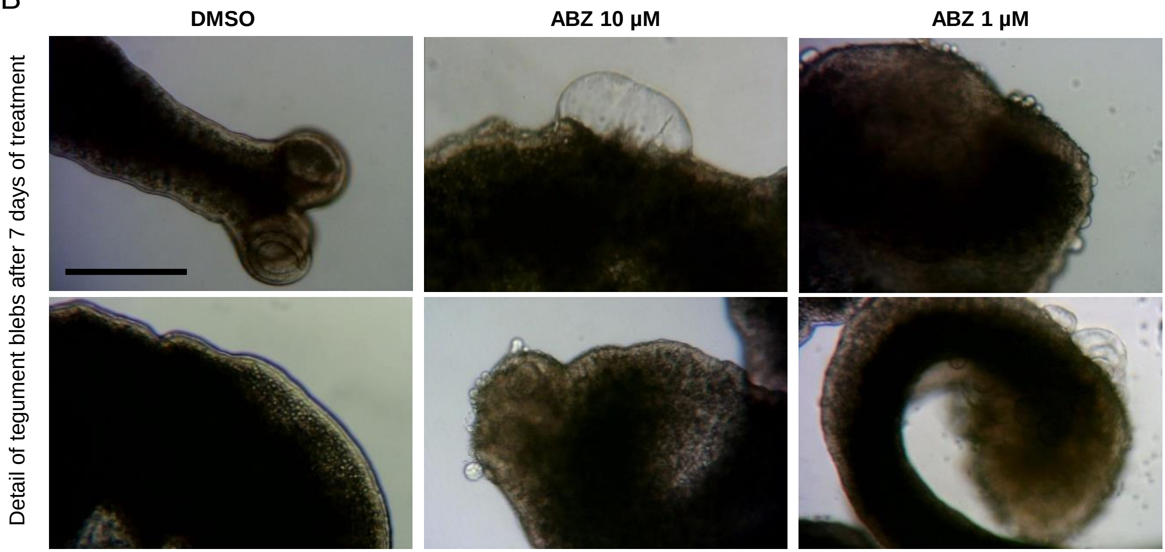

Supplementary Figure 4

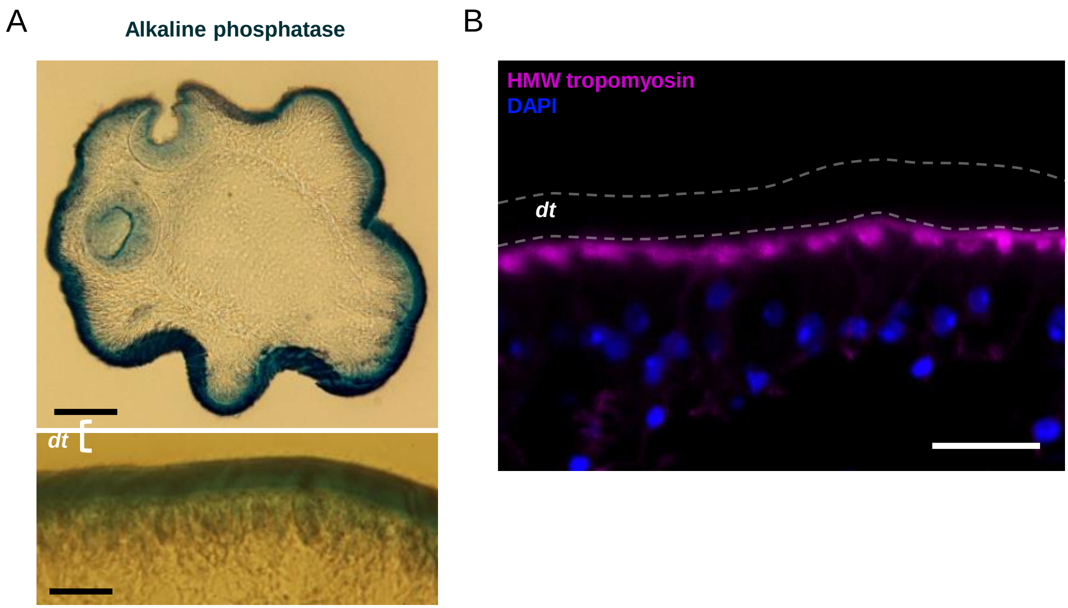

Supplementary Figure 5

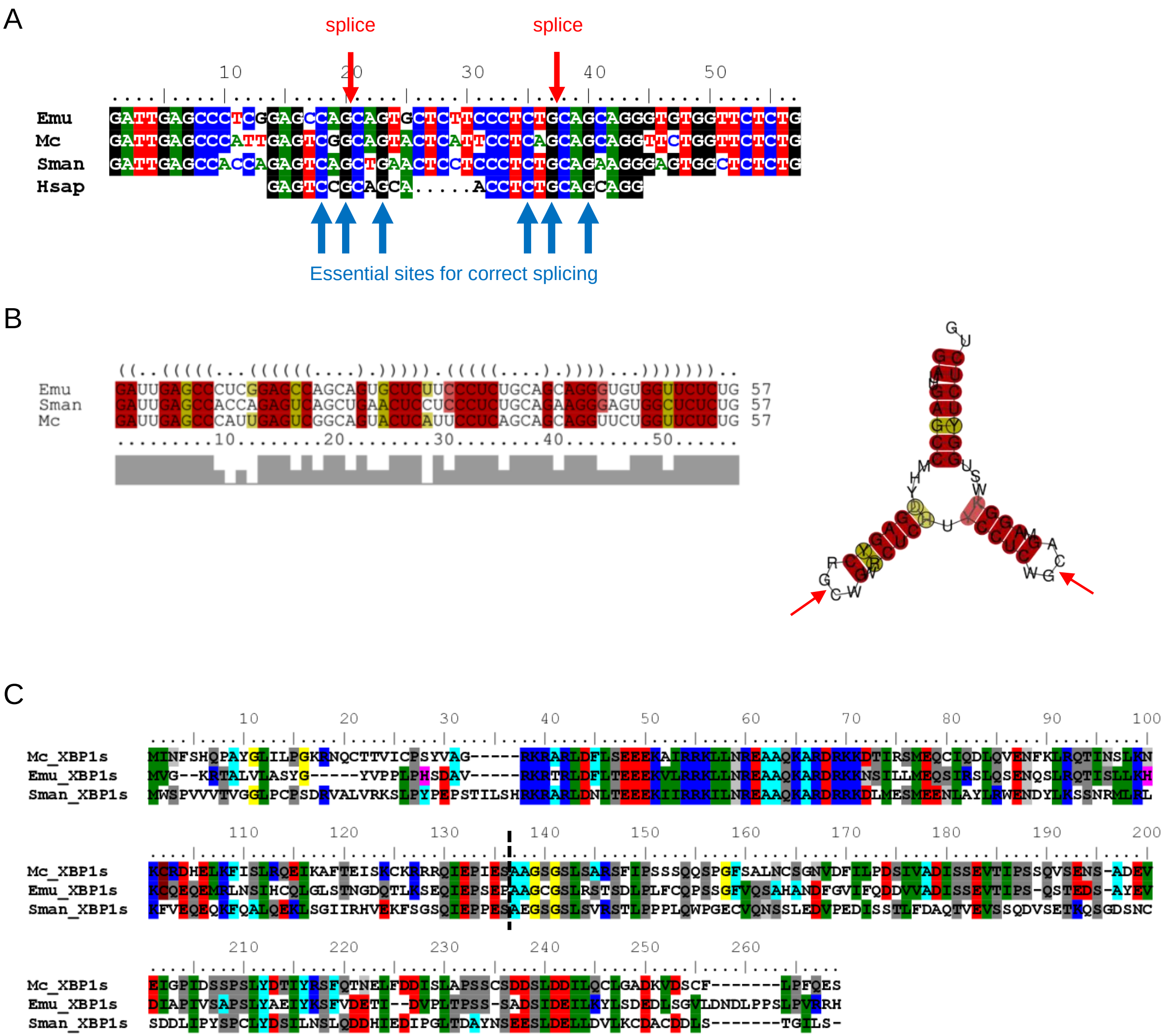

Supplementary Figure 6

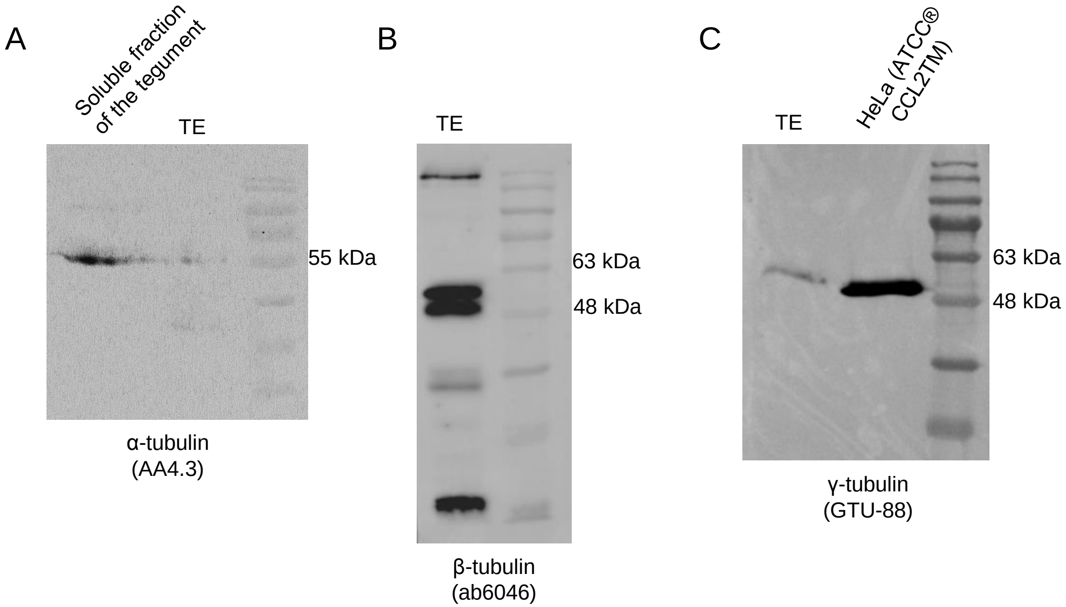

| Supplementary Table 1. Antibodies and their dilutions | | | | |
|---|---|---|---|---|
| **Primary antibodies** | **Dilution for IHF** | **Dilution for Western Blot** | **Original epitope** | ***M. corti* epitope** |
| AA4.3 (DSHB, U.S.A.; anti-α-tubulin, mouse monoclonal, culture supernatant. RRID: AB_579793) | 1 in 10 | 1 in 200 | | |
| GTU-88 (Sigma, T5326, Israel; anti-γ-tubulin, mouse monoclonal. RRID: AB_532292) | 1 in 250 | 1 in 1000 | EEFATEGTDRKDVFFY | EEFATNGSDRKDVFFY |
| ab6046 (Abcam, U.S.A.; anti-β-tubulin, rabbit polyclonal. RRID: AB_2210370) | 1 in 200 | 1 in 500 | | |
| 6-11B-1 (Sigma, T7451, U.S.A.; anti-acetylated tubulin, mouse monoclonal. RRID: AB_609894) | 1 in 500 | - | | |
| Y10b (DSHB, U.S.A.; anti-rRNA, mouse monoclonal, culture supernatant. RRID: AB_2313703) | 1 in 3 | - | | |
| 12D10 (Millipore, MABE343, U.S.A; anti-puromycin, mouse monoclonal. RRID: AB_2566826) | 1 in 500 | - | | |
| Anti-High Molecular Weight Tropomyosins (Koziol et al 2011; rabbit polyclonal). | 1 in 500 | 1 in 1000 | | |
| Anti-McU_009411 (GeneScript; anti-Alkaline Phosphatase, rabbit polyclonal against LPEVDLPQPPYPIKC-KLH ) | - | 1 in 1000 | | |
| Anti-Tyrosinated-tubulin (Thermo, MA1-80017, United Kingdom; rat monoclonal. RRID: AB_2210201) | 1 in 100 | - | | |
| Anti-Phospho-eIF2α (Invitrogen, 44-728G, India; rabbit polyclonal. RRID: AB_2533736) | - | 1 in 1000 | SELSRRRIRSINK | SELSRRRIRSINK |
| **Secondary antibodies** | | | | |
| Anti-Mouse-Alexa 555 Highly cross-adsorbed (Thermo, A21424, U.S.A. RRID: AB_141780) | 1 in 500 | - | | |
| Anti-Rabbit-Alexa 546 (Thermo, A11010, U.S.A. RRID: AB_2534077) | 1 in 1000 | - | | |
| Anti-Rat-Alexa 488 Highly cross-adsorbed (Thermo, A48262, U.S.A. RRID: AB_2896330) | 1 in 200 | - | | |
| Anti-Mouse-HRP (Thermo, A16084, U.S.A. RRID: AB_2534758) | - | 1 in 2000 | | |
| Anti-Rabbit-HRP (Thermo, A16104, U.S.A. RRID: AB_2534776) | - | 1 in 2000 | | |
